## Supplemental Figures for "Functional host-specific adaptation of the intestinal microbiome in hominids"

### **Title**

**Supplementary Tables**

All supplementary tables are found in the supplied excel file

**S1**	Sample Overview

**S2**	All metagenome-assembled genomes and representatives from the UHGGv2 and Manara et al.; including genome score (Magscot score), taxonomic annotation

**S3**	Abundances of all SGBs across the human and African great ape samples

**S4**	P-values from the phylosymbiosis analysis

**S5**	Results from the analysis of group-specific enrichments for microbial genera

**S6**	Results from the analysis of group-specific enrichments for microbial functions

**S7**	Results from the meta-analysis of group-specific enrichments for KEGG annotations

**S8**	Family level results of enrichment analysis of microbial functions differing between humans living in Europe or Africa

**S9**	Meta-analysis results of enrichment analysis of microbial functions differing between humans living in Europe or Africa

**S10**	Enrichment analysis of KEGG annotations differing between humans living in Europe or Africa

**S11**	Results of the enrichment analysis of microbial functions (KOs) differing between NHAs and humans within microbial genera

**S12**	Meta-analysis of the enrichment analysis of microbial functions (KOs) differing between NHAs and humans within microbial genera

**S13**	Results of the co-phylogeny analysis of 209 microbial subtrees

**S14**	Enrichment and depletion of cophylogeny signals for 26 microbial families with SGBs included in the cophylogeny analysis

**S15**	Information on genome size, gene count and 62 in silico inferred traits for 1017 SGBs included in the cophylogeny analysis

**S16**	Results for the 45 traits subjected to the phylogenetic mixed model analysis

Supplementary Figures

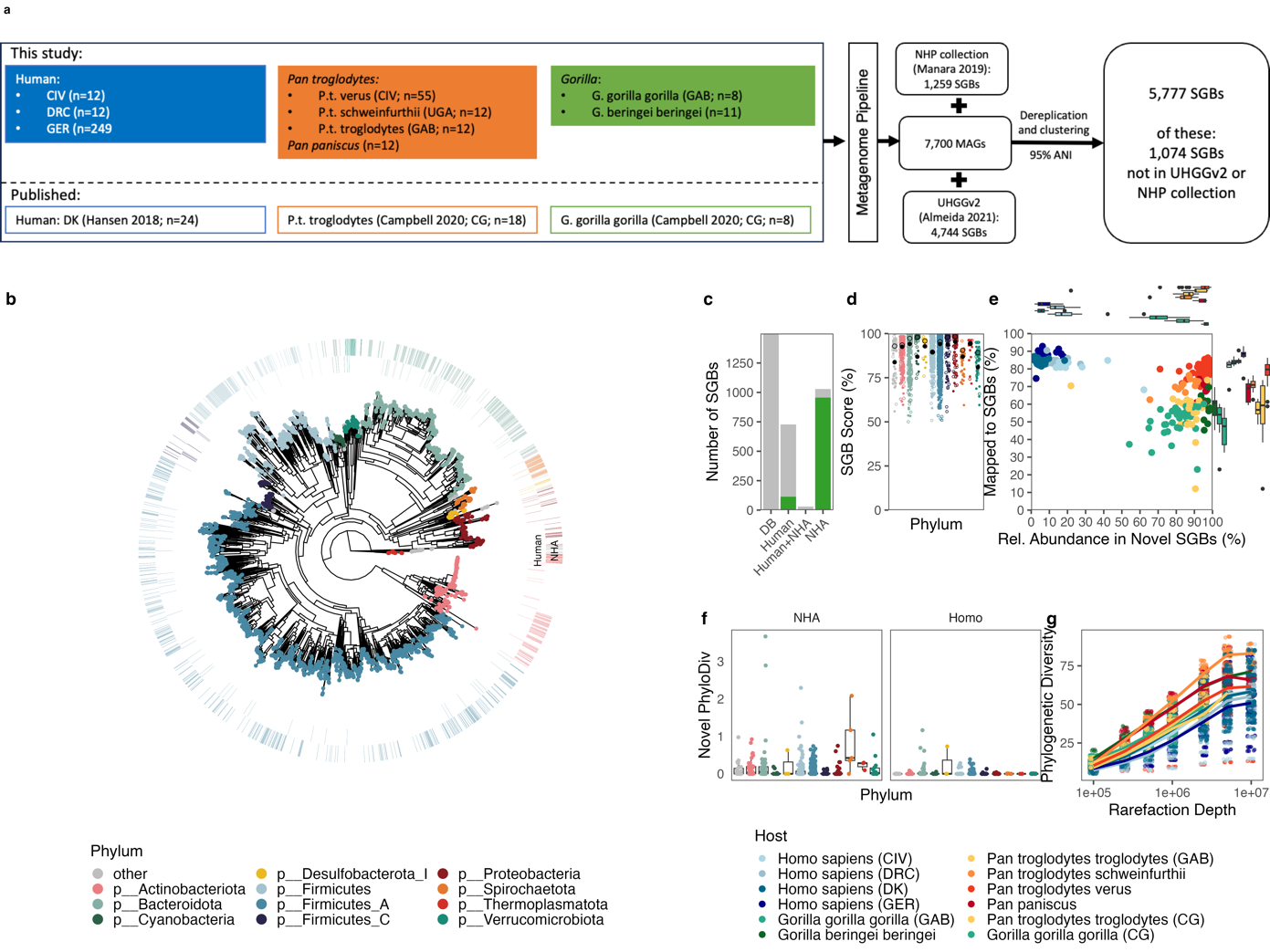

**Supplementary Figure 1:** Genome statistics of newly reconstructed MAGs and representative MAGs from UHGGv2 and Manara et al. (2019). (a) overview on included datasets and data processing (b) Phylogeny of all 2,943 SGB representative MAGs based on 120 bacterial and 53 archaeal universal single-copy marker genes. (c) SGB origin and novelty (green) based on reconstructed MAG sequences from human and NHP hosts, as well as reference databases (UHGGv2 and Manara et al. 2019). (d) MAGScoT scores of all SGB representative genome sequences. Filled shapes represent previously recovered SGBs, filled shapes represent novel SGBs. (e) Mapping success and relative abundance in novel SGBs for the different host groups. (f) overall novelty by NHP and Human host groups measured by Faith’s phylogenetic diversity. (g) per-sample and per-group phylogenetic diversity in relation to metagenomic rarefaction depth.

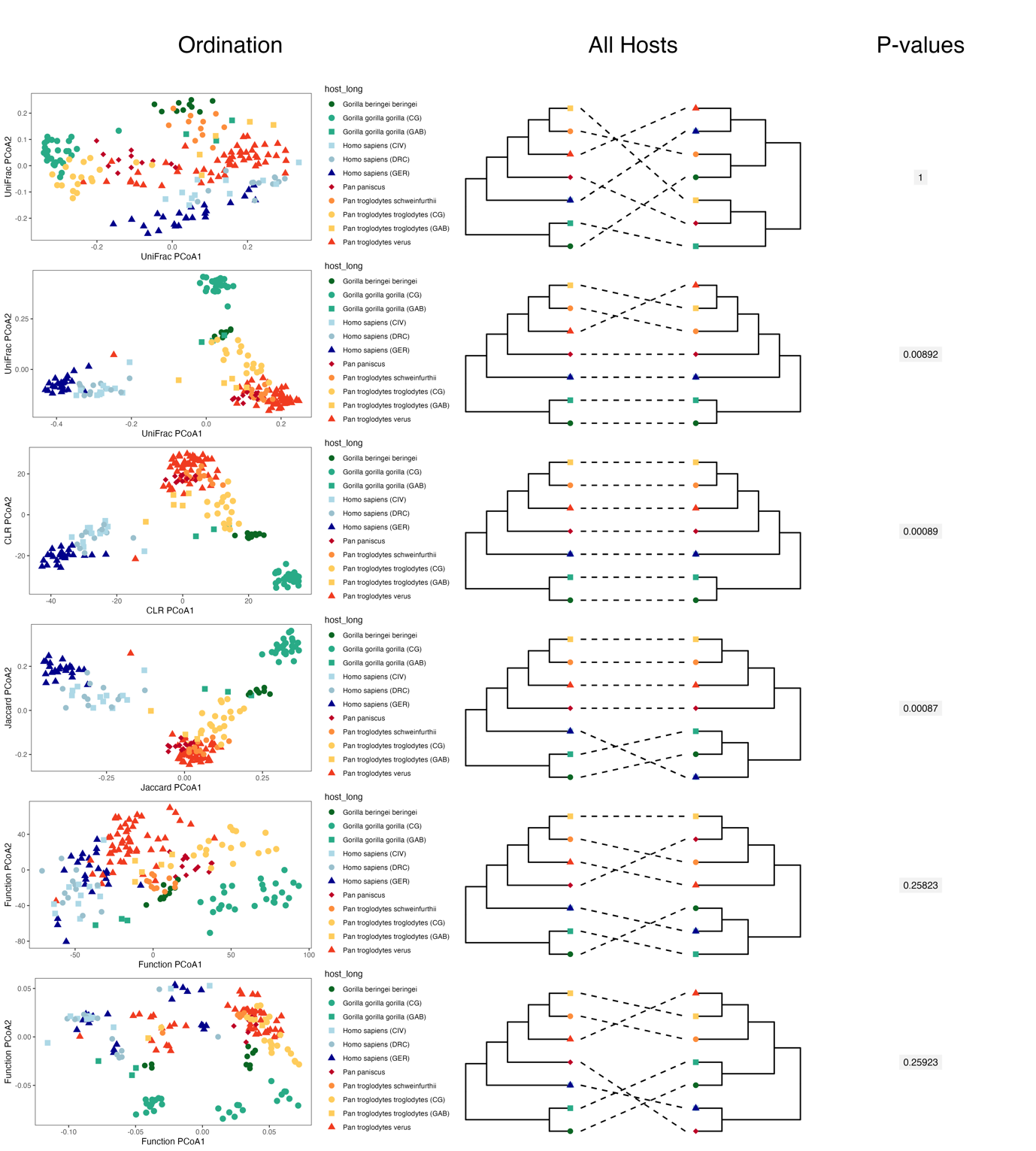

**Supplementary Figure 2:** Ordination, hierarchical clustering and results from the phylosymbiosis analysis for six distance measures: weighted UniFrac, unweighted UniFrac, genus-level Aitchison distance (CLR-transformed genus abundances), genus-level Jaccard distance, as well as distances based on KEGG ontology functional abundances and presence/absence patterns of KEGG ontology functions.
